# Gene replacement therapy provides benefit in an adult mouse model of Leigh syndrome

**DOI:** 10.1101/2020.01.08.894881

**Authors:** Robin Reynaud Dulaurier, Giorgia Benegiamo, Elena Marrocco, Racha Al-Tannir, Enrico Maria Surace, Johan Auwerx, Michael Decressac

## Abstract

Mutations in nuclear-encoded mitochondrial genes are responsible for a broad spectrum of disorders among which Leigh syndrome (LS) is the most common in infancy. No effective therapies are available for this severe disease mainly because of the limited capabilities of the standard adeno-associated viral (AAV) vectors to transduce both peripheral organs and the central nervous system (CNS) when injected systemically in adults.

Here, we used the brain-penetrating AAV-PHP.B vector to reinstate gene expression in the *Ndufs4* KO mouse model of LS. Intravenous delivery of an AAV.PHP.B-Ndufs4 vector in 1-month old KO mice restored mitochondrial complex I activity in several organs including the CNS. This gene replacement strategy extended lifespan, rescued metabolic parameters, provided behavioral improvement, and corrected the pathological phenotype in the brain, retina, and heart of *Ndufs4* KO mice. These results provide a robust proof that gene therapy strategies targeting multiple organs can rescue fatal neurometabolic disorders with CNS involvement.

## Introduction

Mitochondrial diseases are a heterogenous group of genetic human disorders characterized by a primary defect in the mitochondrial oxidative phosphorylation system. About a third of these diseases results from a deficiency in complex I activity and represents a wide spectrum of clinical manifestations ranging from fatal neonatal diseases to adult-onset neurodegenerative pathologies (Kirby *et al.*, 1999). Leigh syndrome (LS), also known as subacute necrotizing encephalomyelopathy, is the most common infantile mitochondrial disease. This fatal pediatric condition is characterized by gliosis, demyelination and necrotic lesions in multiple brain areas including the brainstem and the basal ganglia that appear as hyperintense signal seen at the MRI. Patients exhibit a variety of symptoms such as psychomotor arrest or decline, ataxia, dystonia, lethargy, abnormal ocular movements and respiratory failure (Kirby *et al.*, 1999). To date, no therapeutic strategy alleviates the pathology and passing often occurs within the decade following diagnosis.

Several mutations have been associated to LS including some in the nuclear gene *Ndufs4* that encodes for a non-enzymatic 18kDa subunit of complex I (Vinothkumar *et al.*, 2014). *Ndufs4* knockout (KO) mice closely replicate the pathological hallmarks observed in LS patients. While *Ndufs4* heterozygous mice appear similar to wild-type animals, KO mice progressively develop encephalopathy leading to an early death around postnatal day (P)50 (Kruse *et al.*, 2008). Starting from P40, mice manifest LS-like symptoms including growth retardation, hypothermia, ataxia, lethargy and breathing irregularities (Kruse *et al.*, 2008). Recent studies also demonstrated the contribution of peripheral organ dysfunction in the phenotypic manifestation highlighting the need for a broad correction of the genetic defect (Quintana *et al.*, 2010; Jin *et al.*, 2014).

At the pre-clinical level, strategies based either on a pharmacological inhibition of mTOR or exposures to hypoxic conditions have provided remarkable therapeutic effects as well as important insights into the disease mechanism but their transposability in the clinic is limited (Johnson *et al.*, 2013; Johnson *et al.*, 2015; Jain *et al.*, 2016; Ferrari *et al.*, 2017). Gene replacement therapy using an AAV9 vector has also been attempted in *Ndufs4* KO neonates but resulted in marginal benefits (Quintana *et al.*, 2012; Di Meo *et al.*, 2017).

In the present study we took advantage of the recent development of a novel AAV9 vector variant able to cross the blood brain barrier also in adult mice (Deverman *et al.*, 2016). We therefore used the brain-penetrating AAV-PHP vector for a gene replacement strategy in *Ndufs4* KO mice and we showed that this approach afforded a long-term therapeutic effect by correcting LS-associated pathology in several organs and resulted in lifespan extension.

## Material and methods

### Animals

*Ndufs4* heterozygous mice were obtained from Jackson Laboratories (stock number 27058) and bred to produce *Ndufs4* KO offsprings. Mice were genotyped at 2 weeks of age using the protocol from Jackson Laboratories and the following primers: wild type forward: AGTCAGCAACATTTTGGCAGT, common: GAGCTTGCCTAGGAGGAGGT, mutant forward: AGGGGACTGGACTAACAGCA. In order to provide warmth and social interaction, KO mice were housed with a minimum of one control littermate. Water bottles had long tip and food was provided on the bottom of each cage containing KO mice so that access to food and water was not a limiting factor for survival. Mice were observed daily and euthanized if they showed a 20% loss in maximum body weight, lethargic behavior, or were found prostate or unconscious. Animals weight was recorded twice a week and body temperature was measured on the same day using a rectal thermometer (Microtherma2, Thermoworks). Measurements were done between 2.00pm and 4.00pm to minimize the effect of circadian variations on body temperature. All experimental procedures were conducted in accordance with the guidelines of the European Directives (2010/63/EU) and the ARRIVE guidelines. They were reviewed and approved by the Italian Ministry of Health, Department of Public Health, Animal Health, Nutrition and Food Safety, in accordance with the law on animal experimentation (article7; D.L. 116/92; protocol number: 26/2014).

### Adeno-associated viral (AAV) vector production and injection

The plasmid containing the *Ndufs4*-IRES-GFP cassette under the control of the CMV-chicken β-actin (CBA) promoter was provided by Pr. Palmiter (University of Washington School of Medicine) (Quintana *et al.*, 2012). The AAV.PHP.B plasmid was provided by Dr. Gradinaru (California Institute of Technology) (Deverman *et al.*, 2016). The AAV-PHP.B-*Ndufs4* and the control vector AAV-PHP.B-GFP vector were produced by the AAV Vector Core (Telethon Institute of Genetics and Medicine (TIGEM) as previously described (Colella *et al.*, 2014).

At 4 weeks of age, KO mice and control littermates were randomly assigned to receive an intravenous injection of the *Ndufs4*-expressing or the GFP control AAV-PHP.B vector (1 × 10^12^ gc, diluted in phosphate buffer saline (PBS)) through the retro-orbital sinus.

### Paw clasping test, cylinder test, hanging test, and seizure frequency

In order to assess neurological decline, the paw clasping test was performed at 45 and 250 days of age. As previously described (Decressac *et al.*, 2010), mice were suspended by the tail and their clasping behavior was examined during 30 seconds.

For the cylinder test, mice were placed in a 2 liters glass cylinder, a video was recorded for 5 min and analyzed using the EthoVision XT software (version 14.0). The number of rears (wall rears, seated rears or free rears; see Fig. 2c and Supplementary videos 1-3) (Schweizer *et al.*, 2016) was scored and the time spend immobile was measured as an index of the locomotor activity.

**Fig. 1:**
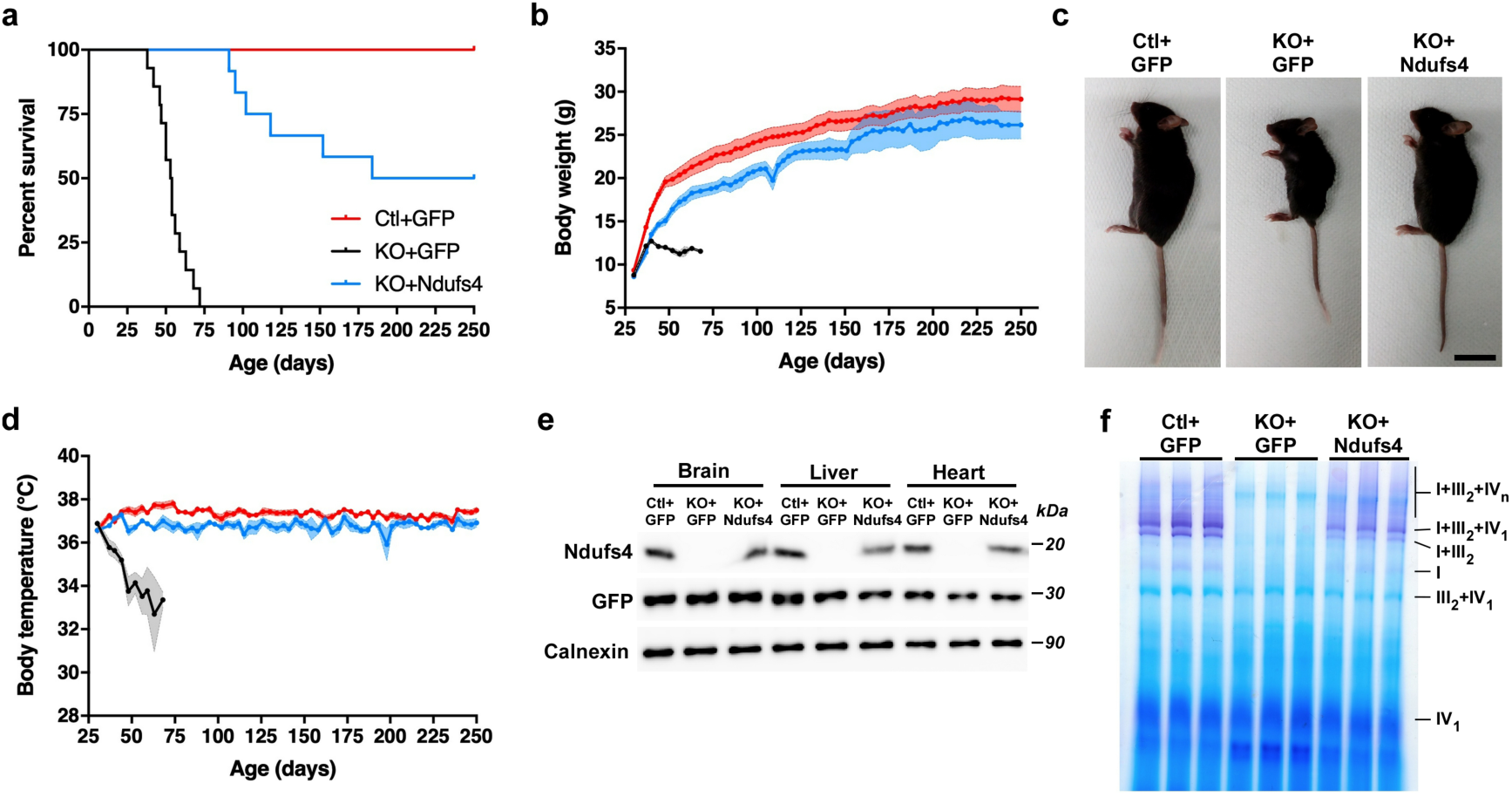
Gene replacement therapy in Ndufs4 KO mice extends lifespan by restoring mitochondrial function. (a) Kaplan-Meier survival curve for control mice injected with the AAV-GFP vector (Ctl+GFP, red curve, n=11) and KO mice injected with the AAV-GFP vector (KO+GFP, black curve, n=14) or with the AAV-*Ndufs4* vector (KO+*Ndufs4*, blue curve, n=12) (log-rank P<0.001 (Mantel-Cox test). (b) Body weight curves starting from the day of AAV vector injection (1 month old) and measured twice a week over the survival period. (c) Representative pictures of 45 days-old mice from the Ctl+GFP, KO+GFP, and KO+*Ndufs4* groups. Scale bar = 1cm). (d) Body temperatures curves starting from the day of AAV vector injection (1-month-old) and measured twice a week over the survival period. (e) Western blotting analysis of Ndufs4, GFP and calnexin in the brain, liver, and heart in 45 days-old mice. (f) BN-PAGE and Complex I in-gel activity assay in isolated brain mitochondria (blue: Coomassie, purple: complex I activity). Individual complexes and super-complexes are labeled. Data in b and d are presented as means ± SEM.

**Fig. 2:**
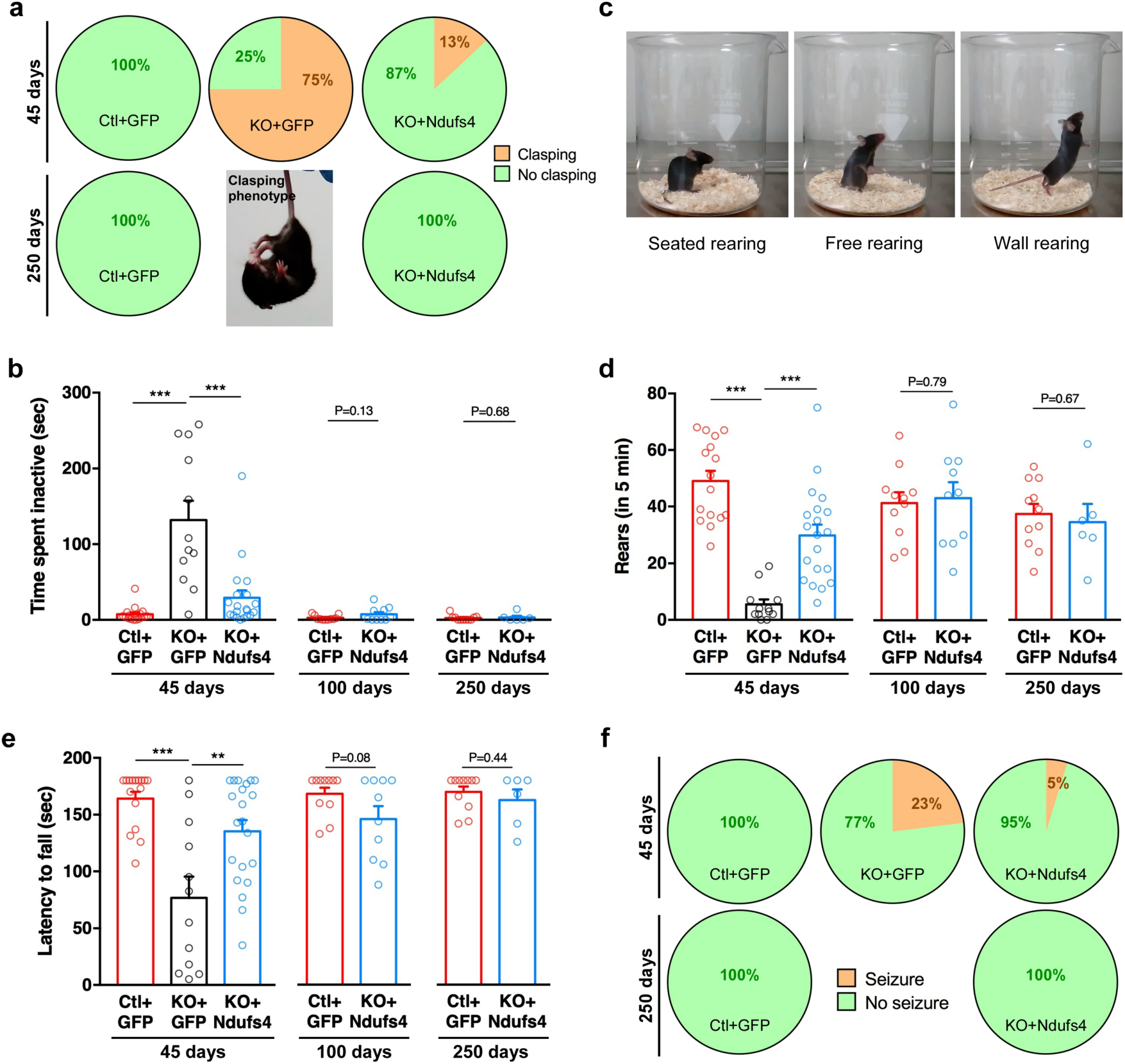
Ndufs4 gene therapy ameliorates behavioral performances. (a) Neurological defect was assessed by the paw clasping test. Pie charts represent the proportion of mice eliciting clasping behavior at 45 and 250 days of age. Insert shows an example of clasping behavior in a KO mouse treated with the AAV-GFP vector. (b-c) General locomotor activity was assessed using the cylinder test. The total time spent immobile (b) and the number of rears (c-d) were recorded during 5 minutes. Representative photographs illustrating the different types of rears that were quantified (c). Data were collected at 45 days of age for all experimental groups (n=12-20/group) (***P<0.001, one-way ANOVA and Tukey’s multiple comparison) and at 100 and 250 days of age for AAV-GFP-treated control mice and AAV-*Ndufs4*-treated KO mice (n=6–11/group) (student’s *t*-test). (e) Strength and resistance was assessed using the hanging test. Data were collected at 45 days of age for all experimental groups (n=12-20/group) (**P<0.01, ***P<0.001 one-way ANOVA and Tukey’s multiple comparison) and at 100 and 250 days of age for AAV-GFP-treated control mice and AAV-*Ndufs4*-treated KO mice (n=6–11/group) (student’s *t*-test). (f) Pie charts representing the proportion of animals exhibiting epileptic seizures at 45 and 250 days of age. Data in b, d, and e are presented as means ± SEM.

The hanging test was performed at 45, 100 and 250 days. Mice were placed on the top of a standard wire cage lid. The lid was lightly shaken to cause the animals to grip the wires then turned upside down and suspended at a height of 40 cm over an open cage filled with bedding and excess nesting materials to prevent injury from falling. The latency for the mice to fall off the wire grid was measured and average values were computed from two trials (15 min apart). Trials were stopped if the mouse remained on the lid after 3 min (Luk *et al.*, 2012).

The percentage of animals manifesting epileptic seizure (Supplementary video 4) was estimated at 45, 100 and 250 days. Mice were monitored for the occurrence of epileptic episodes during 3 hours. At the end of an epileptic event, the identity of the mouse was recorded and the animal returned to its home cage.

All testing and analysis were performed by an experimenter blind to the phenotype and treatment.

### Electroretinogram (ERG)

The method used for electrophysiological recordings (ERGs) was described previously (Botta *et al.*, 2016). Briefly, mice were dark-adapted for 3 hours. After anaesthesia, mice were accommodated in a stereotaxic apparatus under dim red light, their pupils dilated with a drop of 1% tropicamide (Alcon Laboratories, Inc., Fort Worth, TX) and the body temperature maintained at 37.5°C. ERGs were evoked by flashes of different light intensities ranging from –4 to +1.3 log cd.s/m^2^, in scotopic condition, generated through a Ganzfeld stimulator (CSO, Florence, Italy). For each animal group, representative a-wave and b-wave track (evoked at + 1.3 cd.s/m^2^ light stimulus) are shown.

### Blood collection and serum lactate analysis

Blood was collected from the retro-orbital plexus and serum was isolated by centrifugation. Lactate concentration was determined using a colorimetric assay following the manufacturer’s instruction (Abcam, ab65331). Measures of absorbance were done on a PHERAstar FS plate reader (BMG Labtech) and collected with the MARS data analysis software version 3.20 R2 (BMG Labtech). Hematocrit was determined by measuring the space occupied by red blood cells after centrifugation of blood in heparinized capillaries (Brand Micro-hematocrit capillaries 749311) at 10,000 rpm for 5 minutes.

### BN-PAGE, total OxPhos immunostaining and complex I in-gel activity assay

The BN-PAGE and in gel activity assay protocols were described in detail previously(Jha *et al.*, 2016). Briefly, ∼30 mg of frozen brain tissue were homogenized in ice-cold isolation buffer (0.2M sucrose, 10mM Tris, 1mM EGTA/Tris pH 7.4, adjust pH to 7.4 with 1M HEPES buffer, protease inhibitors). Following centrifugation, isolated mitochondria protein content was quantified using DC protein assay (Bio-rad). For BN-PAGE immunoblotting and in-gel activity, 50μg of mitochondria extract were solubilized using 5% digitonin. Electrophoresis of solubilized mitochondrial proteins was performed using the NativePAGE system (Novex) using 3-12% gradient gels. For immunoblotting samples were run at 150V for 30 min and at 250V for additional 90min. Proteins were transferred on a PVDF membrane using an iBlot Gel Transfer device (Invitrogen) and incubated with primary antibodies (Anti-OxPhos Complex Kit (Life technologies, cat. no. 457999) and Anti-MTCO1 antibody (Abcam, ab14705) to detect total OxPhos proteins. Immunostaining of the membrane was performed using Western Breeze Chromogenic Immunodetection System (Invitrogen). For complex I in-gel activity assay samples were run at 150V for 30 min and at 250V for additional 150min to obtain maximal separation of supercomplexes bands. Gels were run at 4°C to preserve enzymatic activity. After electrophoresis gels were incubated with Complex I substrate solution (2mM Tris-HCl, 0.1 mg/ml NADH, 2.5 mg/ml Nitrotetrazolium Blue chloride) at RT until the appearance of purple bands indicative of complex I activity (10-20 min).

### Tissue staining and microscopy

Mice were deeply anesthetized by intra-peritoneal injection of ketamine and were then perfused through the ascending aorta with PBS. Tissues were dissected and post-fixed during 24 hours at 4**°**C in 4% paraformaldehyde. Tissues were cryoprotected overnight in 25% sucrose before being embedded in OCT and cut on a cryostat (Leica CM3050S or Thermo Fisher Scientific CryoStar NX50).

Immunohistological staining was performed as previously described(Decressac *et al.*, 2012). Sections were washed with PBS, incubated in 0.1 M PBS containing 10% normal goat serum, 0.25% Triton X-100 and the primary antibodies for 16–18 hours at room temperature, shaking gently. The tissue sections were then washed three times in PBS and incubated in secondary antibodies secondary antibodies coupled to Alexa 488 or Alexa 568 (1:300, Molecular Probes, Invitrogen) diluted in the blocking solution for 2 hours at room temperature. Sections were rinsed again and coverslipped using the Vectashield Hard Set mounting medium (Vector Laboratories, Burlingame, CA) or the Prolong Gold Antifade medium (Molecular Probes).

Primary antibodies were used to detect GFAP (rabbit, 1:2000, Abcam, #ab7260), Iba1 (rabbit, 1:2000, Wako, #019-19751), GFP (chicken, 1:2000, Abcam, #ab13970), Calbindin (mouse, 1:1000, Millipore, #ab1778), NeuN (rabbit, 1:1000, Cell Signaling, #12943), TH (mouse, 1:3000, Immunostar, #22941).

Staining of F-actin on liver samples was performed by incubating sections with Alexa Fluor 594 Phalloidin (1:600; Molecular Probes, A12381) together with the secondary antibodies.

For the staining of lipid droplets, brain sections were rinsed in PBS and then incubated for 10 min in Nile red (2.5μg/ml dissolved in 75% aqueous glycerol). Subsequently, sections were washed twice with PBS, mounted and coverslipped with Prolong Gold Antifade medium (Molecular Probes). Images were captured using the LSM 710 confocal microscope (Zeiss).

### Quantification of retinal ganglion cells and Purkinje cells

The number of retinal ganglion cells was determined by counting NeuN-positive cells in the retinal ganglion cell layer. The number of Purkinje cells was determined by counting calbindin-positive cells in the cerebellum. Images were analyzed using ImageJ v.2.0.0 software (National Institute of Health, Bethesda, MD, USA). Quantification was performed in 10 random fields of view per mouse and the average number of NeuN-positive or calbindin-positive cells per mm was calculated.

### Analysis of cardiac pathology

Analysis of cardiac pathology was first determined by calculating the heart weight (g) / body weight (g) ratio. In addition, we performed a morphometric analysis of the cardiac tissue. 35μm-thick heart sections were cut on a cryostat (Leica CM3050S or Cryostar NX50) followed by staining with wheat germ agglutinin (WGA) and DAPI. Free floating sections were washed in HBSS and stained with WGA-CF™ 640 (5μg/ml, HBSS) (Biotium) for 4 hours at 37°C to visualize cell membranes. Nuclei were stained with DAPI (5μg/ml, PBS) (Invitrogen) for 5 min before being rinsed 3 times in PBS. Sections were mounted on glass slides and coverslipped with the ProLong Gold Antifade mounting medium (Molecular Probes). Images of entire sections were acquired using the Axio Scan.Z1 slide scanner (Zeiss) and images of the WGA staining were captured with a LSM 710 confocal microscope (Zeiss). For each animal, cardiomyocytes cross-sectional area was measured from 10 different pictures of WGA-stained sections. A minimum of 250 cells were analyzed using ImageJ v.2.0.0 software (National Institute of Health, Bethesda, MD, USA).

### Western blotting

Tissues were harvested and homogenized in RIPA buffer (Sigma). 20 to 40μg of protein were boiled at 95**°**C for 5 minutes in Laemmli buffer (Biorad), separated on a SDS-PAGE gel and then electrotransfered (100V, 1 hour) on a PVDF membrane (Millipore). After blocking for 1 hour in Tris-buffered saline with 0.1% Tween-20 (TBS-T) and 3% non-fat dry milk, membranes were incubated overnight at 4**°**C with one of the following primary antibodies: GFP (chicken, 1:2000, Abcam, #ab290), GFAP (rabbit, 1:2000, Santa-Cruz, #sc-6170), Ndufs4 (mouse, 1:2000, Abcam, #ab87399), Calnexin (Rabbit, 1:5000, Enzo Life Science, ADI-SPA-860). After washing for 30 minutes in TBST, membranes were incubated for 1 hour at room temperature with an HRP-conjugated secondary antibody (1:1000, Promega). Protein expression was revealed using the Clarity kit (Biorad). Luminescence signal was detected using the GE Amersham AI600 (GE Healthcare) or the ChemiDoc MP (Biorad) and band intensities were quantified by densitometry using ImageJ v.2.0.0 software (National Institute of Health, Bethesda, MD, USA).

### Statistical analysis

Statistical analysis was conducted using the GraphPad Prism software (version 7.0a). One-way ANOVA test with Tukey’s multiple comparison test were performed to analyze the difference between experimental groups. Student’s *t*-test was used to analyze the difference between two groups. The data were collected and processed in a randomized and blinded manner. No statistical methods were used to predetermine sample size, but our sample sizes are similar to those generally employed in the field. All values are presented as mean ± standard error of the mean (SEM). Statistical significance was set at P<0.05.

### Data availability

All the data and reagents (if not commercially available) that support the findings of this study are available from the corresponding author upon reasonable request.

## Results

*Ndufs4* KO mice present with severe pathology in several organs including the brain, the retina, the liver and the heart (Kruse *et al.*, 2008; Karamanlidis *et al.*, 2013; Chouchani *et al.*, 2014; Jin *et al.*, 2014; Yu *et al.*, 2015). We first verified that the AAV.PHP.B vector was efficient in transducing these tissues following intravenous injection in 1-month-old mice. Two weeks after AAV vector injection, transgene expression was detectable throughout the central nervous system (Supplementary Fig. 1a). GFP-positive cells were observed in the olfactory bulb, striatum, hippocampus, midbrain, cerebellum, spinal cord and vestibular nucleus and co-localized with ubiquitous or subtypes-specific neuronal markers (Supplementary Fig. 1b-c). The AAV-PHP.B vector was also efficient in inducing transgene expression in other tissues including the liver, heart, muscle and retina (Supplementary Fig. 1c-d).

These results prompted us to examine the effect of systemic delivery of an AAV-PHP.B-*Ndufs4* vector (thereafter referred as AAV-*Ndufs4* vector) in the *Ndufs4* KO mouse model of LS. One-month-old KO and control mice were randomly assigned to receive an intravenous injection of an AAV-*Ndufs4* or a GFP-expressing control vector (1 × 10^12^ genome copies/mouse) and they were regularly monitored throughout their survival period. In line with previous reports, KO mice treated with the control GFP vector died at a median age of 54 days. In contrast, gene replacement therapy substantially increased the lifespan of LS mice as 50% of the KO treated with the AAV-*Ndufs4* vector survived 250 days (log-rank P<0.0001; χ^2^=38.36) (Fig. 1a). Bi-weekly monitoring of the body weight of the animals showed that delivery of the *Ndufs4* gene improved this parameter and promoted a continuous, yet slightly delayed, growth of KO mice (Fig. 1b-c). At 5-6 months of age, AAV-*Ndufs4*-treated KO mice were barely distinguishable from control animals (P=0.23 at 170 days) (Fig. 1b). Before dying, the body temperature of *Ndufs4* KO mice falls progressively below 34°C (Fig. 1d) (Jain *et al.*, 2016). Gene replacement therapy rescued hypothermia as KO animals treated with the AAV-*Ndufs4* vector maintained a core body temperature similar to those of control mice (Fig. 1d) (P>0.05). We confirmed that these physiological improvements were associated with a restoration of *Ndufs4* expression in the brain, liver, heart and muscle (Fig. 1e). Complex I is essential for the assembly of mitochondrial super-complexes (Moreno-Lastres *et al.*, 2012). Consistent with the recovery of *Ndufs4* expression, blue native in-gel activity assay and total OxPhos immunostaining demonstrated the physical restoration of complex I and super-complexes as well as an improvement of mitochondrial complex I activity (Fig. 1e; Supplementary Fig. 2).

As LS primarily affects the brain and causes severe neurological symptoms in patients, we first assessed the paw clasping phenotype in mice (Johnson *et al.*, 2013). While 75% of AAV-GFP treated KO mice displayed clasping behavior at 45 days of age, only 13% showed this phenotype in the AAV-*Ndufs4* treated group (Fig. 2a). Notably, none of the KO mice treated with the AAV-*Ndufs4* vector showed this phenotypic sign at 250 days of age (Fig. 2a). In addition, the cylinder test revealed a long-lasting improvement of locomotor activity in the AAV-*Ndufs4* treated group as seen by a reduction in the time spent inactive and a higher number of rears compared to KO mice treated with the control vector (Fig. 2b-d and Supplementary video 1-3). Likewise, long-term improvement in strength and resistance were observed in the hanging test (Fig. 2e). Brain pathology was also illustrated by the occurrence of epileptic seizures in about 23% of KO mice treated with the GFP-expressing viral vector (Fig. 2f and Supplementary Video 4) (Quintana *et al.*, 2010). In contrast, only 5% of KO mice injected with the AAV-*Ndufs4* vector elicited epileptic episodes (Fig. 2f).

As seen in LS patients, *Ndufs4* KO mice show necrotic plaques and severe neuroinflammation in various brain regions including the cerebellum, the vestibular nucleus and the olfactory bulb (Kruse *et al.*, 2008; Johnson *et al.*, 2013). In line with our observation that the AAV-PHP.B vector is efficient in infecting these brain areas and provides improvement in behavioral performances, western blot and histological analysis of the microglia (Iba1) and astrocyte (GFAP) markers confirmed the presence of severe gliosis in the brain of 45 day-old KO mice and revealed that AAV-mediated gene therapy prevented the development of this inflammatory reaction (Fig. 3a-d). In parallel, lipid accumulation in glial cells resulting from the neuronal metabolic distress was markedly reduced at 45 and 250 days of age in the brain of mice treated with the AAV-*Ndufs4* vector (Fig. 3e) (Liu *et al.*, 2015; Liu *et al.*, 2017). To assess the effect of gene replacement on neurodegeneration, we quantified the number of calbindin-positive Purkinje cells in the cerebellum and we found that AAV-*Ndufs4* vector delivery rescued a large number of these neurons in KO mice at both 45 and 250 days (Fig. 3f-g) (Quintana *et al.*, 2010).

**Fig. 3:**
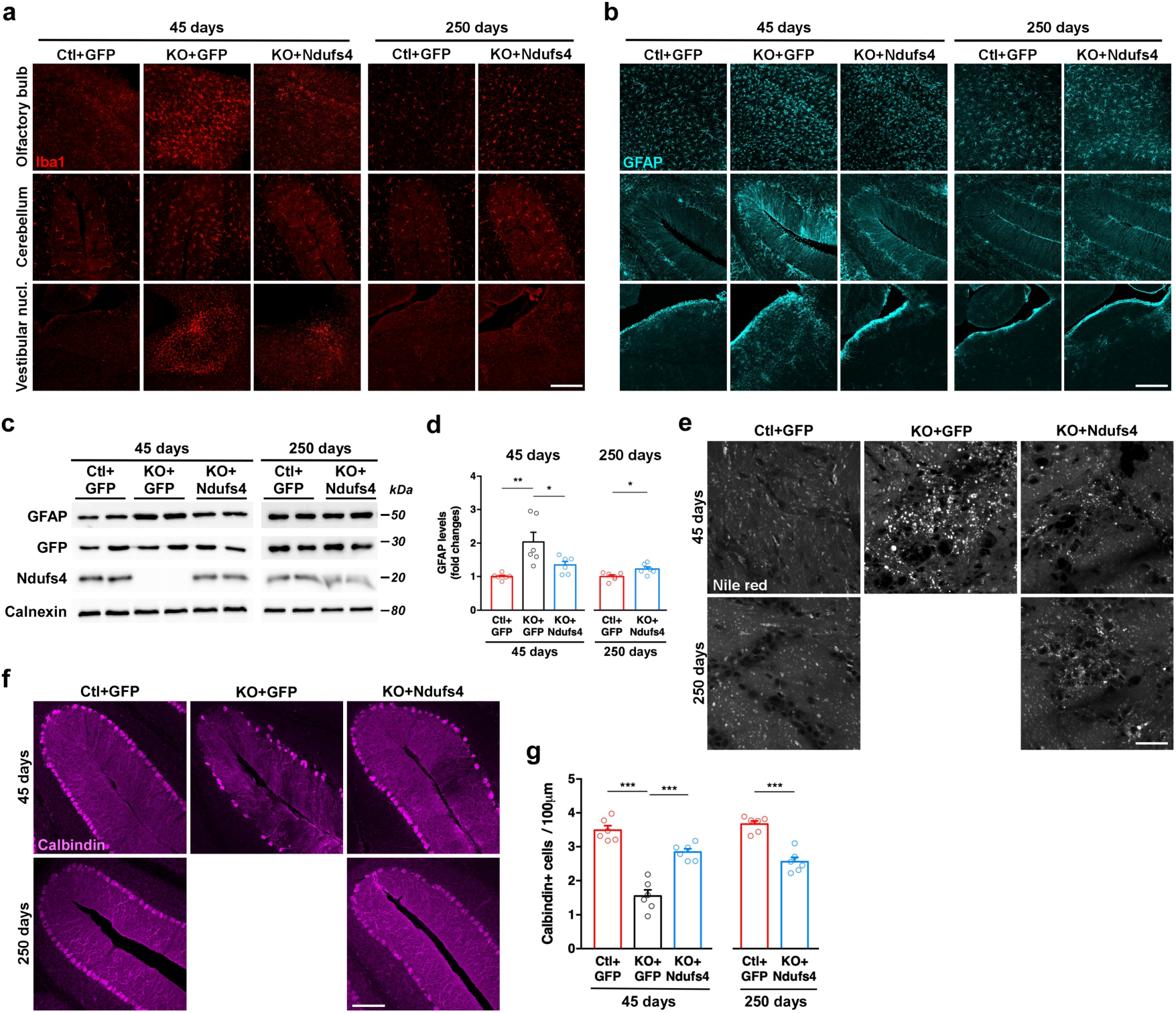
Restoration of Ndufs4 expression in the brain prevented neuronal and glial pathology. (a-b) Representative microscopy images of the olfactory bulb, cerebellum and vestibular nucleus stained for Iba1 (a) or GFAP (b) at 45 and 250 days. Scale bar: 200μm for olfactory bulb and cerebellum, 400μm for vestibular nucleus. (c) Western blots showing the expression level of GFAP, GFP, Ndufs4 and calnexin in the brains of 45-day and 250-day old mice. (d) Quantification of immunoblots (n=6/group) (*P<0.05, **P<0.01, one-way ANOVA with Tukey’s multiple comparison for 45 days results, student’s *t*-test for 250 days results). (e) Lipid accumulation in the brain was detected using Nile red staining. Scale bar: 30μm. (f) Representative microscopy images of calbindin-positive Purkinje cells in the cerebellum at 45 and 250 days. Scale bar: 150μm. (g) Quantification of calbindin-positive neurons in the cerebellum (n=6 per group) (***P<0.001, one-way ANOVA with Tukey’s multiple comparison for 45 days results, student’s *t*-test for 250 days results) Data in d and g are presented as means ± SEM.

In addition to the brain, the retina is severely affected in the LS mouse model (Yu *et al.*, 2015; Song *et al.*, 2017). To test whether restoration of Ndufs4 expression in the retina by gene delivery translated into improved retinal function, we performed electroretinogram recordings and demonstrated that restoration of Ndufs4 expression was associated with the maintenance of the B-wave response that was absent in KO mice (Fig. 4a-b). In line with these results and with our observation that peripheral delivery of the PHP-B vector efficiently transduced the inner retina including the ganglion cell layer and the inner nuclear layer (Supplementary Fig. 1d), we showed that the injection of the AAV-*Ndufs4* vector in *Ndufs4* KO mice protected NeuN-positive retinal ganglion cells as compared to the KO+GFP group (P<0.001) (Fig. 4c-d).

**Fig. 4:**
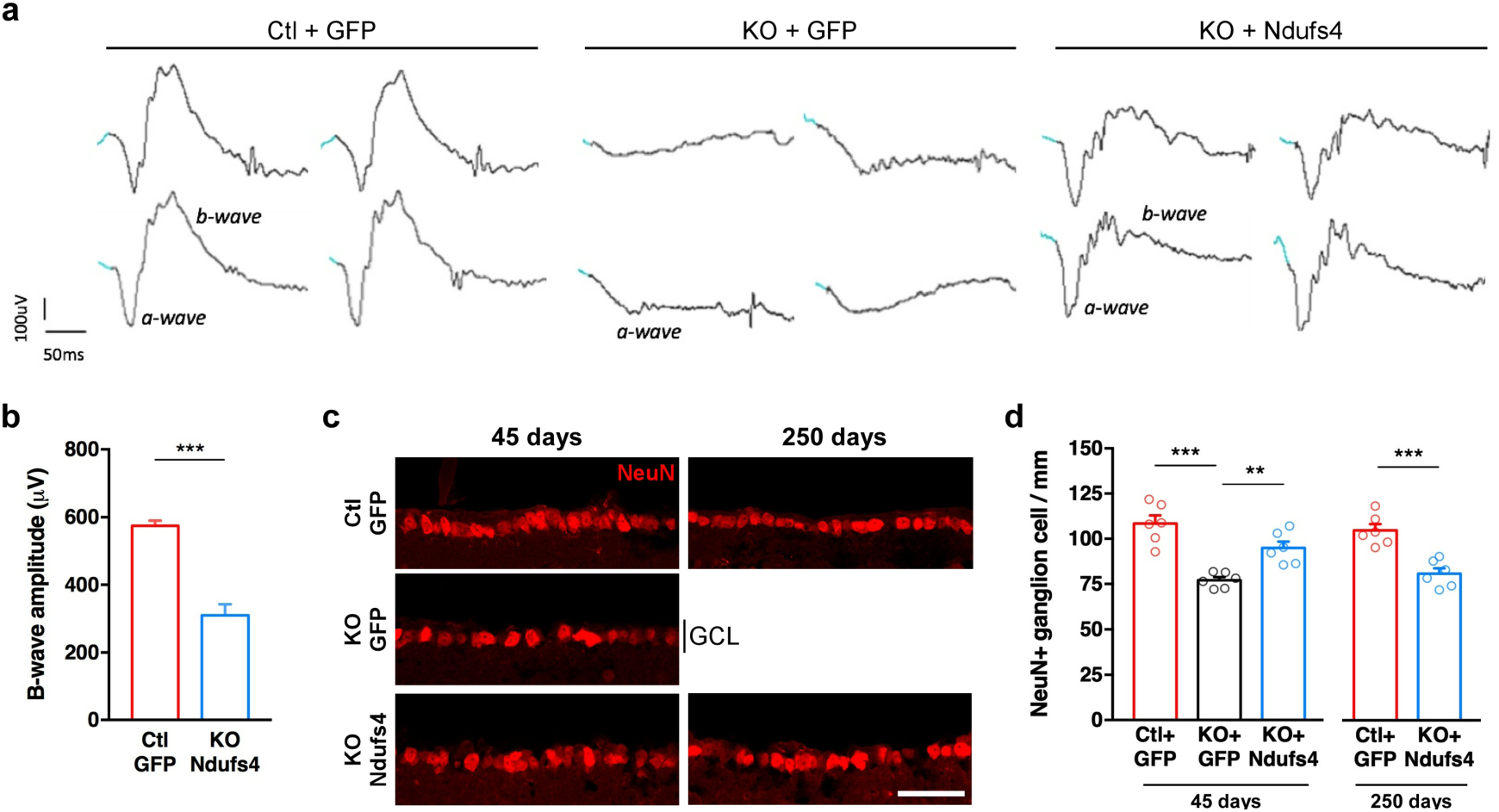
Correction of retinal pathology and retinal function. (a) Representative dark-adapted *in vivo* ERG recordings. Both eyes from two animals are presented for each group. (b) Quantification of B-wave amplitude. Values for the KO+GFP group are not shown as no B-wave was detected. (c) Representative confocal microscopy images from the retina showing NeuN-positive cells from the ganglion cell layer (GCL) in 45 and 250 days-old mice. Scale bar: 50 μm (d) Quantification of NeuN-positive ganglion cells (n=6/group) (**P<0.01,***P<0.001, one-way ANOVA with Tukey’s multiple comparison for 45 days results, student’s *t*-test for 250 days results). Data in b and d are presented as means ± SEM.

Besides the brain pathology, *Ndufs4* KO exhibit peripheral symptoms including metabolic perturbations (Johnson *et al.*, 2013; Jin *et al.*, 2014; Jain *et al.*, 2016). We observed that gene replacement was effective in normalizing the polycythemia (Supplementary Fig. 3a) and in lowering the level of blood lactate in *Ndufs4* KO mice (Supplementary Fig. 3b). In addition, it has been demonstrated that deletion of the *Ndufs4* gene drives hypertrophic cardiomyopathy (Karamanlidis *et al.*, 2013; Chouchani *et al.*, 2014). Consistent with our observation that systemic delivery of the AAV-PHP.B vector efficiently targeted the cardiac tissue (Supplementary Fig. 1c), we showed that treatment with the *Ndufs4*-expressing vector corrected this cardiac abnormality both at the anatomical and cellular level (Supplementary Fig. 3c-f).

## Discussion

Our study provides a proof-of-principle that gene replacement can exert a robust therapeutic effect in an adult mouse model of LS. These results have two important implications. We report the first-ever successful gene therapy with long-term benefits in this pre-clinical model of severe mitochondrial disease. This is an important step considering that previous attempts resulted in marginal extension of lifespan. Indeed, targeted delivery of a *Ndufs4*-expressing AAV vector to the vestibular nuclei or combined systemic and intracerebroventricular injections of a similar viral vector in neonates slightly increased the median survival to 65 and 82 days, respectively (Quintana *et al.*, 2012; Di Meo *et al.*, 2017). In contrast, *Ndufs4* KO mice treated with rapamycin or exposed to hypoxic conditions survived more than 100 days (Johnson *et al.*, 2013; Johnson *et al.*, 2015; Jain *et al.*, 2016; Ferrari *et al.*, 2017). Our results indicate that widespread gene replacement improves survival to a degree at least as good as the ones reported for these non-genetic strategies. Although we performed a single-dose study, one can anticipate that lower doses of AAV.PHP.B vector would afford milder therapeutic effects. This assumption is based on the results from both the AAV-based strategy and the non-genetic approaches which showed that reduced dosage or intermittent treatment have reduced efficacy (Johnson *et al.*, 2013; Johnson *et al.*, 2015; Jain *et al.*, 2016; Di Meo *et al.*, 2017; Ferrari *et al.*, 2017). It is essential to emphasize that gene therapy should remain the ultimate goal for such genetic condition as it is the only strategy that can permanently and efficiently fix the primary defect. Pharmacological or other non-genetic approaches could nevertheless serve as adjunctive treatments to alleviates residual defects.

Regardless of the clinical relevance of the strategy used, it is fundamental to first report that a fatal disease such as LS can be rescued. This goes along with the identification of the organs driving the main pathological manifestation. Despite the conditional ablation of *Ndufs4* in the brain clearly established that the CNS plays a critical role in LS pathology (Kruse *et al.*, 2008; Quintana *et al.*, 2010), the demonstration that reinstating its expression is feasible and provides a therapeutic benefit had to be made. Similarly reports have been important steps for other rare diseases with multi-organ deficiency such as lysosomal diseases or multiple system atrophy and have paved the road for future clinical development (Hua *et al.*, 2011; Spampanato *et al.*, 2011). The clinical transferability of the tools, methods and doses used to make this first demonstration are questions that need to be addressed in future experiments. For our study, the dose (1 × 10^12^) was selected based on previous reports describing widespread and strong transgene expression in mice (Deverman *et al.*, 2016; Challis *et al.*, 2019). Although recent studies have shown that the AAV.PHP.B vector produced a different and more restricted pattern of expression in non-human primates as compared to mice, this new AAV variants remains a powerful tools for initial demonstration of therapeutic rescue as reported here (Hordeaux *et al.*, 2018; Liguore *et al.*, 2019).

The second important information provided by our work is that delayed intervention can still rescued the disease phenotype even if the therapeutic window to intervene is narrow as it is the case for pediatric conditions like LS. Despite the pathology is already prominent in one-month old *Ndufs4* KO mice, we decided to administer the AAV vector at a stage where most organs are almost fully mature hence reducing the risk of a time-dependent decrease in transgene expression (Di Meo *et al.*, 2017). This timeline is also clinically relevant since children are diagnosed after the presentation of symptoms.

Beyond the findings reported here in a mouse model of LS, our study brings promising perspectives for other fatal conditions with CNS involvement as well as for less severe pathologies with later onset.

## Acknowledgments

We thank Edoardo Nusco, Mathilde Faideau, the TIGEM AAV Vector Core, and the TIGEM animal facility staff for excellent technical assistance. We thank the Photonic Imaging Center of Grenoble Institute Neurosciences (Univ. Grenoble Alpes – Inserm U1216) which is part of the ISdV core facility and certified by the IBiSA label. We are grateful to Pr. Richard D. Palmiter (University of Washington, Seattle) for providing the pAAV-CBA-*Ndufs4*-IRES-GFP plasmid and to Dr. Viviana Gradinaru for providing the PHP.B packaging plasmid.

This work was supported by the Fondazione Telethon (grant #TMMDMTX16TT to M.D), by the Agence Nationale de la Recherche (ANR-JCJC program, grant #ANR-17-CE37-0008-01 to M.D), by the grants from the Agence Nationale de la Recherche ANR-15-IDEX-02 NeuroCoG (to M.D) in the framework of the “Investissement d’Avenir” program, by the European Research Council (ERC-AdG-787702 “UPRmt” to J.A, and ERC.311682 “Allelechoker” to E.M.S), by the Swiss National Science Foundation (SNSF 310030B-160318 to J.A), by the Strategic Focal Area “Personalized Health and Related Technologies (PHRT)” of the ETH Domain (grant #2018-422 to G.B), by a GRL grant of the National Research Foundation of Korea (NRF GRL 2017K1A1A2013124 to J.A) and by the EPFL. M.D is supported by an IDEX Chair of Excellence from the University of Grenoble-Alpes and by the Edmond J. Safra Foundation. M.D laboratory is member of the Grenoble center of Excellence in Neurodegeneration. The authors also want to thank the Fondation Bettencourt Schueller and the Plan Cancer.

Authors declare that no competing financial and/or non-financial interests exist.

## Authors contribution

R.R.D, G.B, E.M, R.A-T, and M.D performed the experiments, R.R.D, G.B, E.M, E.M.S and M.D prepared figures, R.R.D, G.B, E.M.S, J.A and M.D designed experiments, R.R.D, G.B, E.M, E.M.S, J.A and M.D wrote the manuscript, R.R.D, G.B, E.M, E.M.S, J.A and M.D analyzed the results.

All authors discussed the results and commented on the manuscript at all stages.

## Supplemental information

**Supplementary Fig. 1:**
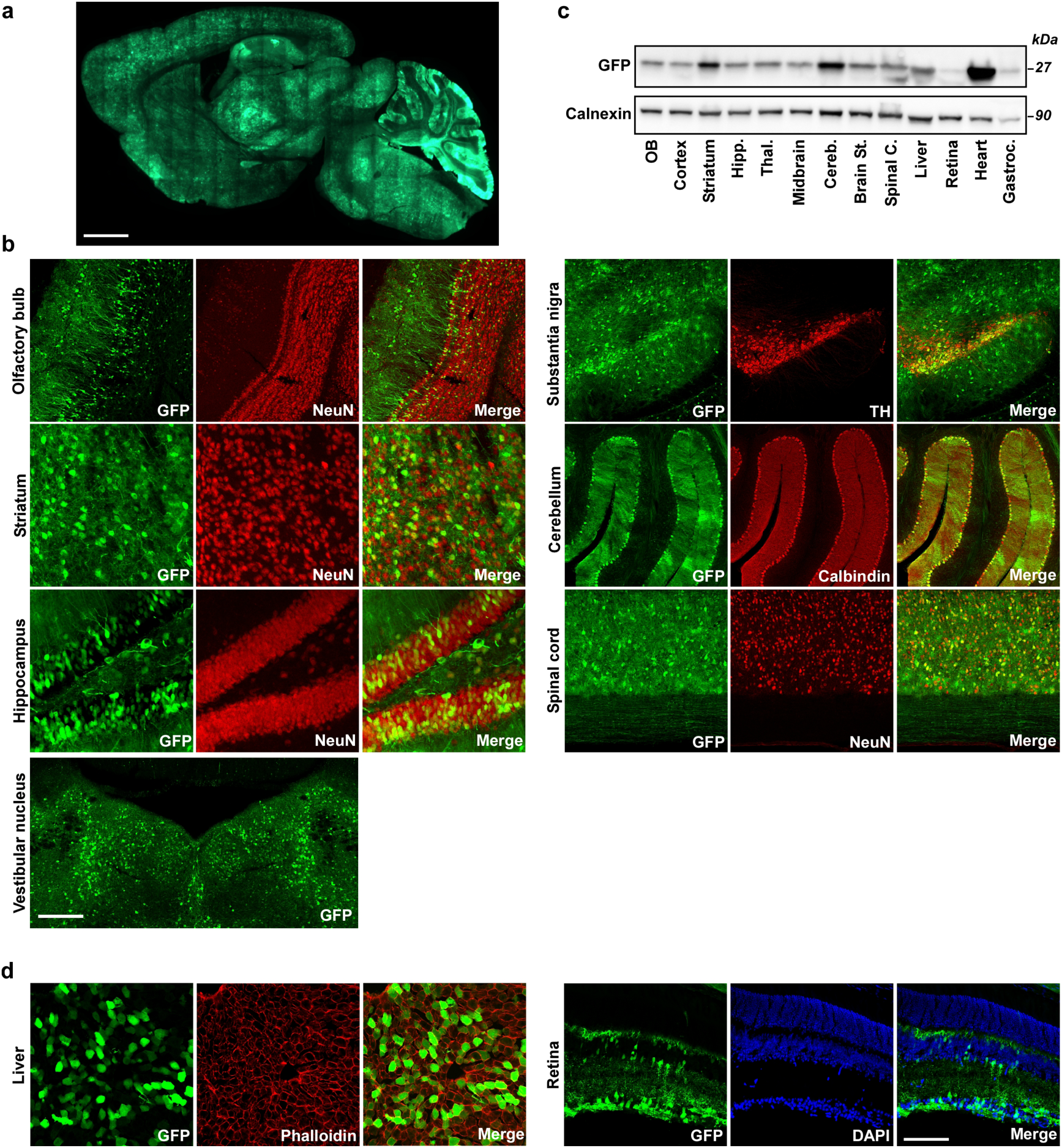
Systemic delivery of the AAV-PHP.B-Ndufs4 vector induces widespread expression of the transgene in the CNS and peripheral organs. (a) Scanning microscope image of a sagittal section from a mouse brain stained for GFP 2 weeks after systemic AAV-PHP.B vector administration. Scale bar: 1.5 mm (b) Confocal microscopy images showing NeuN-positive neurons (in red) in the olfactory bulb, the striatum, the hippocampus, TH-positive (in red) nigral dopamine neurons, and calbindin-positive (in red) Purkinje cells in the cerebellum expressing the transgene (GFP in green) 2 weeks after systemic AAV vector injection. Scale bar: 250 μm for the olfactory bulb, 100 μm for the striatum and hippocampus, 350 μm for the substantia nigra, 400 μm for the cerebellum, 300 μm for the spinal cord and the vestibular nucleus. (c) Western blot analysis showing the expression of GFP in various brain regions, heart liver, muscle, retina 2 weeks after AAV-PHP.B vector injection. (d) Confocal images showing the expression of GFP (in green) in the liver (Phalloidin in red) and in the retina, including retinal ganglion cell layer and inner nuclear layer (DAPI in blue). Scale bar: 120 μm for the liver and the retina.

**Supplementary Fig. 2:**
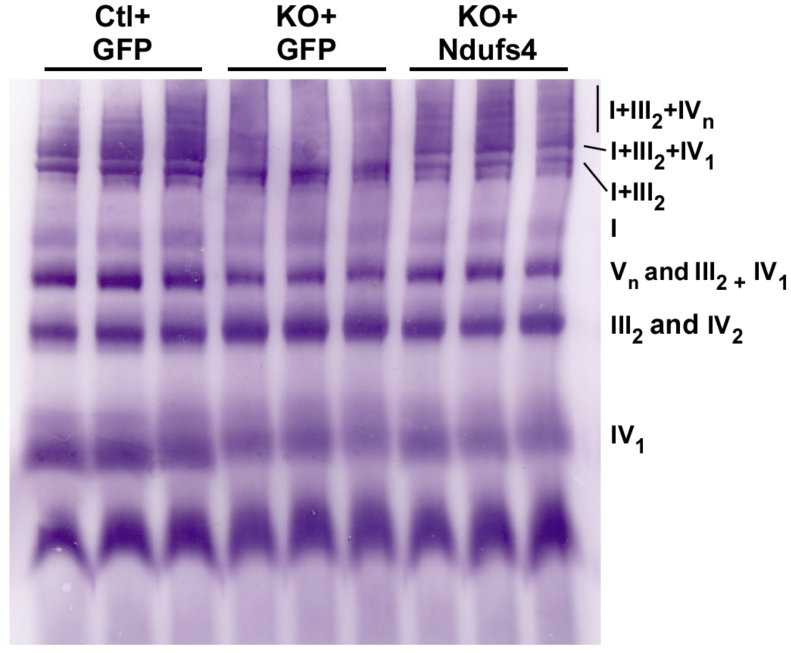
BN-PAGE immunoblot (total OxPhos cocktail) in isolated brain mitochondria. Individual complexes and supercomplexes are labeled.

**Supplementary Fig. 3:**
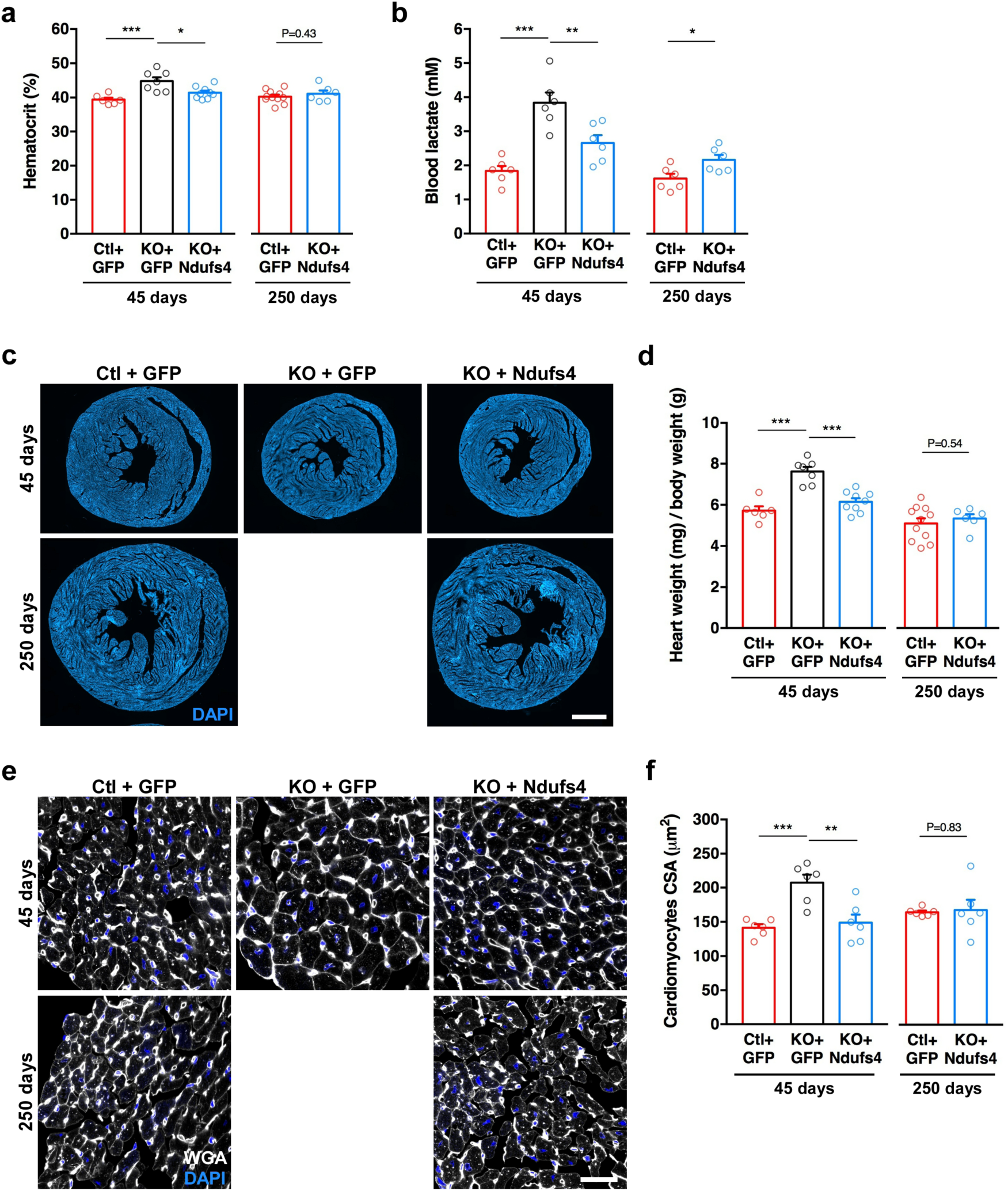
Ndufs4 gene replacement rescued the cardiac pathology. (a) Hematocrit levels were measured at 45 and 250 days of age. (n=6-11/group) (*P<0.05, ***P<0.001; one-way ANOVA with Tukey’s multiple comparison for 45 days results, student’s *t*-test for 250 days results). (b) Blood lactate levels at 45 and 250 days of age. (n=6/group) (**P<0.01, ***P<0.001; one-way ANOVA, Tukey’s multiple comparison for 45 days results, student’s *t*-test for 250 days results). (c) Representative pictures of horizontal heart sections stained with DAPI (blue). Scale bar: 1mm. (d) Cardiac hypertrophy was assessed by calculating the heart weight / body weight ratio of 45-days old mice. (n=6-11/group). (**P<0.01, ***P<0.001; one-way ANOVA, Tukey’s multiple comparison for 45 days results, student’s *t*-test for 250 days results). (e) Representative pictures of horizontal sections from cardiac tissues stained for wheat germ agglutinin (WGA, in white) and DAPI (blue). Scale Bar: 25μm. (f) Measure of cardiomyocytes cross-sectional area (CSA) from WGA-stained sections (n=6/group) (*P<0.05, **P<0.01; one-way ANOVA with Tukey’s multiple comparison for 45 days results, student’s *t*-test for 250 days results). Data in a, b, d, and f are presented as means ± SEM.

***Supplementary Video 1***: Cylinder test performed at 45, 100 and 250 days by the same mouse from the Ctl+GFP group.

***Supplementary Video 2***: Cylinder test performed at 45, 100 and 250 days by the same mouse from the KO+GFP group.

***Supplementary Video 3***: Cylinder test performed at 45, 100 and 250 days by the same mouse from the KO+*Ndufs4* group.

***Supplementary Video 4***: Epilepsy seizure observed in a 45 days-old KO mouse treated with the AAV-PHP.B-GFP vector.

## References

Botta S, Marrocco E, de Prisco N, Curion F, Renda M, Sofia M, et al. Rhodopsin targeted transcriptional silencing by DNA-binding. Elife 2016; 5: e12242.

Challis RC, Ravindra Kumar S, Chan KY, Challis C, Beadle K, Jang MJ, et al. Systemic AAV vectors for widespread and targeted gene delivery in rodents. Nat Protoc 2019; 14(2): 379–414.

Chouchani ET, Methner C, Buonincontri G, Hu CH, Logan A, Sawiak SJ, et al. Complex I deficiency due to selective loss of Ndufs4 in the mouse heart results in severe hypertrophic cardiomyopathy. PLoS One 2014; 9(4): e94157.

Colella P, Trapani I, Cesi G, Sommella A, Manfredi A, Puppo A, et al. Efficient gene delivery to the cone-enriched pig retina by dual AAV vectors. Gene Ther 2014; 21(4): 450–6.

Decressac M, Kadkhodaei B, Mattsson B, Laguna A, Perlmann T, Bjorklund A. alpha-Synuclein-induced down-regulation of Nurr1 disrupts GDNF signaling in nigral dopamine neurons. Sci Transl Med 2012; 4(163): 163ra56.

Decressac M, Wright B, Tyers P, Gaillard A, Barker RA. Neuropeptide Y modifies the disease course in the R6/2 transgenic model of Huntington’s disease. Exp Neurol 2010; 226(1): 24–32.

Deverman BE, Pravdo PL, Simpson BP, Kumar SR, Chan KY, Banerjee A, et al. Cre-dependent selection yields AAV variants for widespread gene transfer to the adult brain. Nat Biotechnol 2016; 34(2): 204–9.

Di Meo I, Marchet S, Lamperti C, Zeviani M, Viscomi C. AAV9-based gene therapy partially ameliorates the clinical phenotype of a mouse model of Leigh syndrome. Gene Ther 2017; 24(10): 661–7.

Ferrari M, Jain IH, Goldberger O, Rezoagli E, Thoonen R, Cheng KH, et al. Hypoxia treatment reverses neurodegenerative disease in a mouse model of Leigh syndrome. Proc Natl Acad Sci U S A 2017; 114(21): E4241–E50.

Hordeaux J, Wang Q, Katz N, Buza EL, Bell P, Wilson JM. The Neurotropic Properties of AAV-PHP.B Are Limited to C57BL/6J Mice. Mol Ther 2018; 26(3): 664–8.

Hua Y, Sahashi K, Rigo F, Hung G, Horev G, Bennett CF, et al. Peripheral SMN restoration is essential for long-term rescue of a severe spinal muscular atrophy mouse model. Nature 2011; 478(7367): 123–6.

Jain IH, Zazzeron L, Goli R, Alexa K, Schatzman-Bone S, Dhillon H, et al. Hypoxia as a therapy for mitochondrial disease. Science 2016; 352(6281): 54–61.

Jha P, Wang X, Auwerx J. Analysis of Mitochondrial Respiratory Chain Supercomplexes Using Blue Native Polyacrylamide Gel Electrophoresis (BN-PAGE). Curr Protoc Mouse Biol 2016; 6(1): 1–14.

Jin Z, Wei W, Yang M, Du Y, Wan Y. Mitochondrial complex I activity suppresses inflammation and enhances bone resorption by shifting macrophage-osteoclast polarization. Cell Metab 2014; 20(3): 483–98.

Johnson SC, Yanos ME, Bitto A, Castanza A, Gagnidze A, Gonzalez B, et al. Dose-dependent effects of mTOR inhibition on weight and mitochondrial disease in mice. Front Genet 2015; 6: 247.

Johnson SC, Yanos ME, Kayser EB, Quintana A, Sangesland M, Castanza A, et al. mTOR inhibition alleviates mitochondrial disease in a mouse model of Leigh syndrome. Science 2013; 342(6165): 1524–8.

Karamanlidis G, Lee CF, Garcia-Menendez L, Kolwicz SC, Jr., Suthammarak W, Gong G, et al. Mitochondrial complex I deficiency increases protein acetylation and accelerates heart failure. Cell Metab 2013; 18(2): 239–50.

Kirby DM, Crawford M, Cleary MA, Dahl HH, Dennett X, Thorburn DR. Respiratory chain complex I deficiency: an underdiagnosed energy generation disorder. Neurology 1999; 52(6): 1255–64.

Kruse SE, Watt WC, Marcinek DJ, Kapur RP, Schenkman KA, Palmiter RD. Mice with mitochondrial complex I deficiency develop a fatal encephalomyopathy. Cell Metab 2008; 7(4): 312–20.

Liguore WA, Domire JS, Button D, Wang Y, Dufour BD, Srinivasan S, et al. AAV-PHP.B Administration Results in a Differential Pattern of CNS Biodistribution in Non-human Primates Compared with Mice. Mol Ther 2019.

Liu L, MacKenzie KR, Putluri N, Maletic-Savatic M, Bellen HJ. The Glia-Neuron Lactate Shuttle and Elevated ROS Promote Lipid Synthesis in Neurons and Lipid Droplet Accumulation in Glia via APOE/D. Cell Metab 2017; 26(5): 719–37 e6.

Liu L, Zhang K, Sandoval H, Yamamoto S, Jaiswal M, Sanz E, et al. Glial lipid droplets and ROS induced by mitochondrial defects promote neurodegeneration. Cell 2015; 160(1-2): 177–90.

Luk KC, Kehm V, Carroll J, Zhang B, O’Brien P, Trojanowski JQ, et al. Pathological alpha-synuclein transmission initiates Parkinson-like neurodegeneration in nontransgenic mice. Science 2012; 338(6109): 949–53.

Moreno-Lastres D, Fontanesi F, Garcia-Consuegra I, Martin MA, Arenas J, Barrientos A, et al. Mitochondrial complex I plays an essential role in human respirasome assembly. Cell Metab 2012; 15(3): 324–35.

Quintana A, Kruse SE, Kapur RP, Sanz E, Palmiter RD. Complex I deficiency due to loss of Ndufs4 in the brain results in progressive encephalopathy resembling Leigh syndrome. Proc Natl Acad Sci U S A 2010; 107(24): 10996–1001.

Quintana A, Zanella S, Koch H, Kruse SE, Lee D, Ramirez JM, et al. Fatal breathing dysfunction in a mouse model of Leigh syndrome. J Clin Invest 2012; 122(7): 2359–68.

Schweizer N, Viereckel T, Smith-Anttila CJ, Nordenankar K, Arvidsson E, Mahmoudi S, et al. Reduced Vglut2/Slc17a6 Gene Expression Levels throughout the Mouse Subthalamic Nucleus Cause Cell Loss and Structural Disorganization Followed by Increased Motor Activity and Decreased Sugar Consumption. eNeuro 2016; 3(5).

Song L, Yu A, Murray K, Cortopassi G. Bipolar cell reduction precedes retinal ganglion neuron loss in a complex 1 knockout mouse model. Brain Res 2017; 1657: 232–44.

Spampanato C, De Leonibus E, Dama P, Gargiulo A, Fraldi A, Sorrentino NC, et al. Efficacy of a combined intracerebral and systemic gene delivery approach for the treatment of a severe lysosomal storage disorder. Mol Ther 2011; 19(5): 860–9.

Vinothkumar KR, Zhu J, Hirst J. Architecture of mammalian respiratory complex I. Nature 2014; 515(7525): 80–4.

Yu AK, Song L, Murray KD, van der List D, Sun C, Shen Y, et al. Mitochondrial complex I deficiency leads to inflammation and retinal ganglion cell death in the Ndufs4 mouse. Hum Mol Genet 2015; 24(10): 2848–60.

